# The Effects of Garlic Extract on Renal Histopathology and Function in a Cadmium Chloride-Induced Hypertension Model

**DOI:** 10.1101/2025.11.19.689172

**Authors:** Gideon B Ojo, Abosede A Olorunnisola, Lawrence D Adedayo, Amos O Morakinyo, Ramotul R Fakunle, Ehidiame S Dawodu

## Abstract

This study investigated the antihypertensive effects of Allium sativum (garlic) on kidney function in a rat model of hypertension.

Forty-eight Wistar rats were divided into groups: control, hypertensive, and hypertensive rats treated with different doses of garlic extract or a standard antihypertensive drug. Hypertension was induced by CaCl2 administration.

The results demonstrated that garlic extract effectively lowered blood pressure in hypertensive rats, indicating its potential as an antihypertensive agent

Further Analysis involve the histopathological examination of kidney tissues and analysis of serum electrolyte levels provided further insights into the renoprotective effects of garlic extract.

This study demonstrates that garlic extract possesses antihypertensive properties in a rat model. Garlic treatment significantly lowered blood pressure compared to the hypertensive control group. Furthermore, kidney tissue analysis revealed potential renoprotective effects. These findings suggest that garlic may offer a promising complementary approach to managing hypertension.

**Graphical Abstract:** 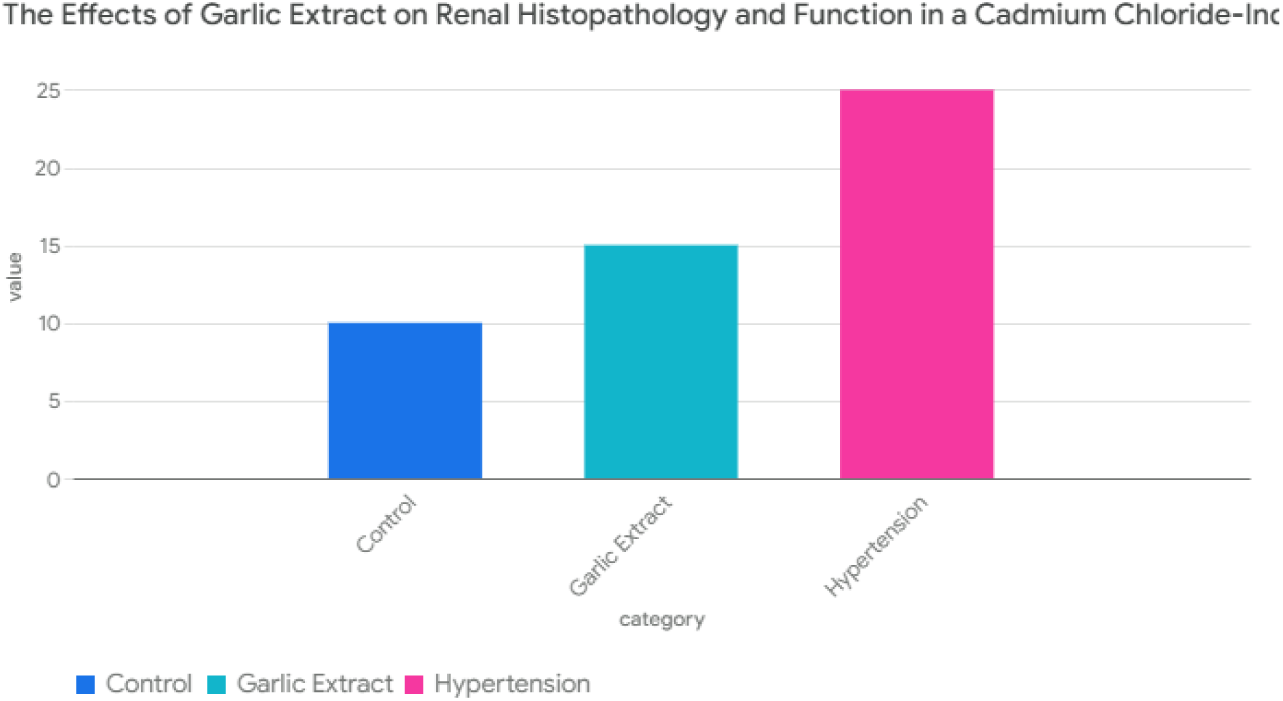

## Introduction

Hypertension, also known as high blood pressure, is a chronic condition characterized by persistently elevated blood pressure within the arteries. While often asymptomatic, it significantly increases the risk of cardiovascular diseases, including stroke, heart attack, and kidney disease. (Poulter *et al.,* 2015).

Typically, hypertension is marked by an increasing trend in systemic arterial pressure that rises above a set benchmark number. The risk of cardiovascular disease (CV) is elevated by raised blood pressure (BP) above 115/75 mm Hg, which grows in a log pattern (Giles et al., 2005).

Among the important organs that are frequently under stress from hypertension and ultimately deteriorate as a result are the kidneys, the eyes, and the heart. High blood pressure, particularly lethal in African-Americans, is associated with almost 75% of particularly lethal in African-Americans and is associated with approximately 75% of all strokes and cardiac arrests. Other risk factors, significant blood pressure elevation, old age, smoking, abnormal cholesterol, family history of early heart disease, obesity, diabetes, coronary artery disease, and signs of another vascular disease are all factors that increase the risk of complications or rapid progression of hypertension (Bellamy et al., 2007).

Primary (essential) hypertension and secondary hypertension are the two types of high blood pressure. The etiology is inherited and non-specific features of lifestyle in 90 to 95 percent of cases. Excessive salt consumption, obesity, smoking, and alcohol usage are all examples of lifestyle decisions that may increase the risk. A further 5-10% of instances are classified as secondary, meaning they have high blood pressure for no apparent reason but may be connected with conditions such as chronic renal disease, arteriosclerosis, endocrine disorders, or the use of birth control pills (Poulter et al., 2015).

Blood pressure is expressed using systolic and diastolic pressures, which have maximum and minimum pressures, respectively. At rest, a person’s blood pressure should be between 60 and 90 mmHg diastolic and 100 to 140 mmHg systolic. The majority of persons have high blood pressure when their resting blood pressure is consistently at or above 140/90 mmHg. Children have different numbers. Poulter et al. (2015) found that ambulatory blood pressure monitoring is more accurate than working blood pressure monitoring. The development of high blood pressure often takes years, and it affects almost everyone. Fortunately, it can be quickly found. According to Kaplan and Victor (2014), uncontrolled high blood pressure increases the chance of significant health issues like heart attack and stroke.

Lower blood pressure reduces the risk of health problems, which can be achieved through medication and lifestyle modifications. Losing weight, eating less salt, exercising more, and adopting a healthy diet are all examples of lifestyle improvements. As a method of treatment, insufficient lifestyle modifications are combined with blood pressure drugs (Poulter et al., 2015).

The global prevalence of hypertension is substantial, with about 1.28 billion adults aged 30–79 years worldwide affected, most of whom live in low- and middle-income countries. The overall global prevalence among adults is around 31%, higher in low- and middle-income countries (31.5%) than in high-income countries (28.5%). (WHO, 2023; NAS, 2025) In specific populations, the prevalence varies. For example: In Nigeria, the age-adjusted prevalence increased to about 32.5% by 2020. (Odili & Chori, 2020)

In the United States during 2021–2023, approximately 47.7% of adults had hypertension, with higher prevalence in men (50.8%) compared to women (44.6%). (CDC, 2024)

In some regions in Nigeria, prevalence rates range widely from about 20.9% in the North-Central region up to 52.8% in the South-East region. (Adeloye *et al.,* 2021)

Prevalence increases with age and higher body mass index (BMI). For instance, hypertension prevalence rises from about 6.8% in people 30 years and younger to over 63% in those older than 70 years, and from 29.2% in lean individuals (BMI <25) up to 75.1% in individuals with BMI ≥40. (Adeloye *et al.,* 2021) Overall, hypertension remains highly prevalent and a leading contributor to cardiovascular disease worldwide, with considerable variations by region, age, and sex. Awareness, treatment, and control rates, especially in low- and middle-income countries, tend to be lower than desirable. (Odili & Chori, 2020; WHO, 2023; NAS, 2025) The kidneys play a primary role in regulating blood pressure by controlling sodium balance through the pressure-natriuresis mechanism. Hypertension often results when this mechanism shifts due to kidney abnormalities, causing impaired sodium excretion and volume overload. This leads to increased systemic vascular resistance and blood pressure. Key pathways involve activation of the renin-angiotensin-aldosterone system (RAAS) and sympathetic nervous system (SNS), which increase renal sodium reabsorption and vasoconstriction. Renal vascular damage from hypertension further diminishes kidney function and exacerbates high blood pressure, creating a vicious cycle. Chronic kidney disease and hypertension are mutually reinforcing, with renal pathology both causing and being worsened by hypertension. (Salem 2002; Taler, 2019; Ameer, 2022; Bia and Galarza, 2024)

Garlic, commonly referred to as *Allium sativum,* is a member of the onion genus. Onion, shallot, leek, chive, and rakkyo are close cousins of this plant (Beaumont et al., 2013; Block, 2010). Garlic has reportedly been used by humans for about 7,000 years and is believed to be a native of Central Asia (Ensminger, 1994). It is a staple in the Mediterranean region and is often used as a condiment across Asia, Africa, and Europe. It has been utilized for both traditional medicine and food flavorings since the time of the Ancient Egyptians (Simonetti, 1990).

Garlic contains 0.1–0.36% volatile oil, which is usually thought to be the source of the majority of garlic’s pharmacological effects. At least 33 sulfur compounds are found in garlic, including aliin, allicin, ajoene, allyl propyl, diallyl, trisulfide, S-allyl cysteine, vinyldithiines, and S-allylmercaptocystein are just a few of the compounds found in allicin. Garlic includes arginine, other amino acids, and their glycosides, in addition to sulfur compounds. Minerals like selenium, and also enzymes such as myrosinase, allinase, and other peroxidases.. More sulfur compounds are found in garlic than in any other Allium species. The therapeutic benefits of garlic as well as its strong odor are both caused by sulfur compounds. The enzyme allinase reacts with the sulfur molecule alliin to produce the smell. Heat deactivates this enzyme, which explains why cooked garlic doesn’t have nearly the same physiological effects as raw garlic or as potent of an odor (Christopher, 2007).

According to Capraz et al. (2006), garlic has mostly gained popularity as a supplementary therapy for lowering blood pressure. In a recent in vitro study, Benavides et al. (2007) examined the vasoactive properties of garlic sulfur compounds that are transformed by red blood cells into hydrogen sulfide, a recognized endogenous cardio-protective vascular cell signaling molecule. In one pilot trial, a high dose of garlic (2400 mg, containing 31.2 mg allicin) was utilized, and overall blood pressures reduced at about 5 hours after the intake, including a drop in diastolic pressure of 16 mm (McMahon and Vargas, 1993).

Garlic is suggested to lower high blood pressure through increasing the synthesis of nitric oxide and hydrogen sulfide, both of which aid in blood vessel relaxation (Hope, 2013).

A 2400 mg garlic pill containing 31.2 mg allicin reduced diastolic BP by 16 mmHg after 5 hours of treatment (McMahon and Vargas, 1993). According to a meta-analysis of 415 patients’ data, diastolic pressure was also lowered by 7.7 mmHg (Silagy and Neil, 1994).

Humans have traditionally utilized Allium sativum, or garlic, as a food and medicine (Lawson, 1998). Garlic’s capacity to reduce blood pressure has been linked to its hydrogen sulfide generation, allicin content from alliin, and the enzyme allinase (Banerjee et al., 2003; Benavide et al., 2007; and Higdon & Lawson, 2005), angiotensin II inhibitory and vasodilatory effects in animal and human cell investigations (Al-Qattan et al. Garlic can reduce blood pressure by relaxing and expanding blood arteries (Healthline, 2005).

High blood pressure, sometimes known as hypertension, is a primary cause of cardiac problems. It is one of the main factors in death and disability from heart disease, kidney failure, heart attack, and stroke. Through the actions of its sulfur compounds and its capacity to decrease fatty substances like cholesterol found in the bloodstream, garlic can help lower blood pressure. Garlic consumption can also help restore low blood pressure to normal (Murray and Michael, 1995). Garlic’s ability to significantly reduce the risk of developing hypertension has been attributed to the presence of an active substance known as garlic sulphides and allicin. Allicin aids in blood vessel relaxation while also lowering blood pressure and harm to the blood. Additionally, it prevents the angiotensin I enzyme’s ability to raise blood pressure by contracting smooth muscles. It greatly aids in lowering cholesterol levels and platelet aggregation thanks to its capacity to degrade fibrinolytic activity in the blood (Godiyal, 2013). The effects of garlic extract on renal histopathology and function in a cadmium chloride-induced hypertension model have been extensively studied, revealing significant protective properties. Garlic, particularly its active compounds like diallyl disulfide (DADS) and diallyltetrasulfide (DTS), demonstrates antioxidant and anti-inflammatory effects that mitigate cadmium-induced renal damage. The following sections outline the key findings from the research.

Protective Mechanisms of Garlic Extract

Antioxidant Activity: Garlic extracts enhance antioxidant enzyme levels, reducing oxidative stress markers such as malondialdehyde (MDA) and increasing glutathione (GSH) levels(Alruhaimi et al., 2024)(Awadallah et al., 2011).

Inflammation Reduction: Garlic administration decreases pro-inflammatory cytokines (e.g., IL-6, TNF-α) and histopathological changes associated with cadmium toxicity (Alruhaimi et al., 2024) (Hashem, 2009).

Histopathological Improvements: Histological examinations show reduced kidney damage, including less collagen deposition and cellular infiltration, in garlic-treated groups compared to controls (Hashem, 2009) (Cha, 1987).

Impact on Renal Function

Biochemical Markers: Garlic extract significantly lowers serum creatinine and urea levels, indicating improved renal function (Hashem, 2009) (Raharjo et al., 2020).

Dose-Dependent Effects: Higher doses of garlic extract correlate with better renal outcomes, with optimal doses identified in various studies (Raharjo et al., 2020)(Cha, 1987).

While garlic extract shows promise in protecting against cadmium-induced renal damage, it is essential to consider that its efficacy may vary based on dosage and the specific context of exposure. Further research is needed to fully understand the mechanisms and potential clinical applications of garlic in heavy metal toxicity.

Garlic’s capacity to promote hydrogen sulfide and nitric oxide synthase synthesis is another factor that lowers blood pressure. According to reports, this is what causes blood vessels to relax. In addition to lowering blood pressure, garlic serves other purposes. In addition to its ability to decrease cholesterol, it has the ability to strengthen the immune system and digestive system of a person. However, because it thins the blood as well and may interact poorly with some supplements and medications, it is advised that everyone seek the advice of a healthcare professional before taking it (Godiyal, 2013).

Amlodipine has been shown to block calcium ions from entering vascular smooth and cardiac muscle cells. Extracellular calcium ion transfer into these cells through specific ion channels via gap junctions is required for the heart’s and smooth muscle contractile actions of the heart and smooth muscle in the vascular system. Amlodipine selectively reduces calcium ion influx across cell membranes, with vascular smooth muscle cells being particularly affected. These unfavorable inotropic effects are not detectable in intact animals at therapeutic levels, but they are in vitro. It has been noted that the (-) isomer of the two sterioisomers [R (+), S (-)] is more active than the (+) isomer (Karmoker et al., 2016).

The class of medications known as calcium channel blockers includes amlodipine. Amlodipine increases blood flow by widening blood arteries. Amlodipine has been used to treat coronary artery disease-related disorders such as hypertension, angina, and other chest pains (Multum, 1996).

## Methodology

### Obtaining Plant Material

The bulbs of *Alium sativum* (Garlic) were purchased at Bodija market Ibadan, Oyo state, Nigeria in April, 2016.

### Identification of Plant Material

The plant material was taken to the department of Botany Obafemi Awolowo University, Ife in Osun state Nigeria. It was identified with voucher number IFE- 17508 at Ife herbarium.

### Preparation of Extract

The cloves were peeled and about 2,000g bulbs of fresh garlic was blended into a paste. The total weight of the blended garlic was about 2,404g. The further process of extraction was done through the following steps; Soaking, Filteration and Evaporation.

### Soaking

The blended garlic was soaked in 5litres of methanol for 72hours (3days) and covered using foil paper to allow penetration of methanol into the plant material to obtain the active components.

### Filtration

The filtration process was done on the third day using a sieve, the filtrate was used and the residue was disposed after the filtration process was done.

### Evaporation

The process was done using a rotator evaporator. Some of the filtrate was kept in a conical flask in the water bath. The Temperature of the water bath was set at 78°C which is the boiling point of methanol, and this caused the methanol to evaporate leaving the extract in the conical flask. The evaporated methanol was collected behind the vacuum pump in around bottom flask. Water from the condenser cooled the vacuum pump.

### Preparation of Standard Drug Solution

Amlodipine was obtained from a pharmacy in Ibadan, Oyo state. The tablets were grounded to fine powder and dissolved in normal saline to get the desired concentration (500µg/kg).

### Chemicals Used

The following are the chemicals used in the course of this experiment.

Cadmium chloride, Amlodipine, Diethyl ether, Normal saline, distilled water, Methanol and 10% Neutral Buffered Formalin (NBF).

### Apparatus Used

The apparatus used in the course of this experiment are; Sensitive weighing balance, Beakers, Centrifuge (Centurion Scientific Limited, United Kingdom), Canula, Wooden and Plastic Industrial cages, Dissecting set, Slides, Foil paper, Plain specimen bottle, Microscope, Cotton wool, Paraffin wax, Syringe, Rotatory Evaporator, Lithium Heparinized bottle.

### Housing, Care and Management of Animals

48 Albino Wistar rats with weight range of between 70 -150kg were the experimental animals. They were housed in wooden cages and fed with standard pelleted feed and given water liberally in animal holdings for proper acclimatization for two (2) weeks before the commencement of the experiment in the animal house of the Department of Anatomy, Bowen University, Iwo, Osun State. The rats were fed and given water every morning. The rats’ weight was checked at arrival and at every 1week interval to monitor their growth and development.

### Induction of Hypertension

Hypertension was induced in the animals through injection of freshly prepared solution of cadmium chloride (1mg/kg) in 0.9% w/v saline, intra-peritoneally daily for one week. The dosage given was calculated based on the average weight of each group.

### Animal Grouping

The Animals were randomly distributed into six (6) groups A, B, C, D, E and F with eight (8) animals in each group. The dosages are usually presented as:

Group A: Animals given 0.9% w/v normal saline for 1week. Group B: Animals given cadmium chloride (1mg/kg) for 1week.

Group C: Animals given cadmium chloride (1mg/kg) for 1week followed by administration of the extract @ (100mg/kg) for 1week

Group D: Animals given cadmium chloride (1mg/kg) for 1week followed by administration of the extract@ (250mg/kg) for 1week.

Group E: Animals receiving cadmium chloride (1mg/kg) for 1week followed by administration of Amlodipine @ (500µg/kg) for 1week.

Group F: Animals given the extract@ (250mg/kg) for 1week.

### Animal Sacrifice

The animals were sacrificed 24 hours after the last treatment was given. However, prior to the sacrifice, the following supplies and tools were made available: a centrifuge, sensitive weighing scale, and dissecting set, dissecting board, anesthesia chamber, neutral buffered formalin (NBF) and specimen bottles.

Using Diethyl Ether in the anesthesia chamber, the animals were put to sleep before being sacrificed in accordance with Public Health Service (PHS) guideline from 2002 on the use and care of experimental animals. Each animal’s heart was punctured in order to collect whole blood into sterile, dry test tubes. After centrifuging the test tubes for 30 minutes at 600 rpm, clear, non-hemolyzed serum was collected and stored in the refrigerator for analysis. The organs to be studied were quickly removed and weighed to prevent them from stiffening and then stored in the specimen bottles containing NBF (Fixative).

The weight of the kidneys was recorded and analyzed. The kidneys were fixed in 10% Neutral Buffered Formalin (NBF) and histologically processed. The tissues were stained with Hematoxylin and Eosin (H&E) stain as well as two specific stains, Periodic Acidic Schiff (PAS) stain and Verhoeff-van Gieson (VVG) stain.

### Serum Electrolytes

For Biochemical analysis whole blood was collected by cardiac puncture from each animal into clean dry test tubes. They were then centrifuged for 30 minutes at 600 rpm. Clear non hemolyzed serum was harvested and kept in the fridge before analysis. Trace elements (K, Na and Mg) levels in plasma were determined using Jenway flame photometer (PFP7) and Atomic Absorption Spectrometer (AAS) (PG990).

### Statistical Analysis

The statistical method applied in this study involves expression of the values in form of means and standard error of mean. The difference of mean between each group was analyzed using one way ANOVA in comparison with the control using Duncan Method at a 0.05 significance level.

## Results

### HISTOMORPHOLOGY OF THE KIDNEY

**PLATE 1.**
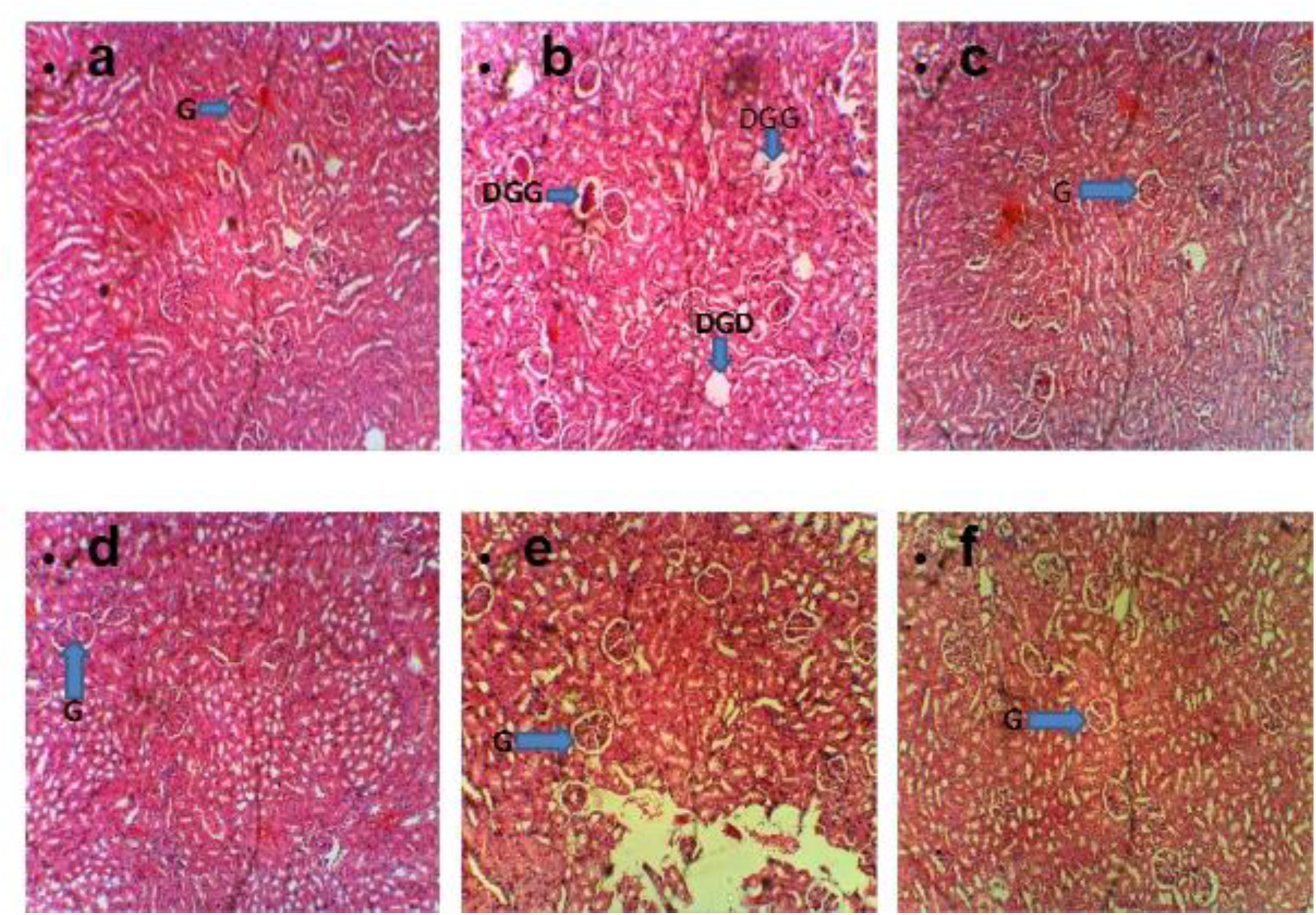
Photomicrograph showing the histology of kidney in Groups A, B, C, D, E and F,using Hematoxylin and Eosin stain (H&E) stain, at X 100 magnification. LEGEND: G – Glomerulus, DGG –Degenerating glomerulus, DGD – Degenerated glomerulus, a – Group A, b – Group B, c - Group C, d - Group D, e – Group E, f – Group F.Group A (control) shows the normal histology of renal parenchyma.Group B (Untreated) shows degenerating or shrinkage glomeruli (DGG) and a degenerated glomerulus (DGD).Group C (CdCl2 + 100mg/kg Extract), Group D (CdCl2 + 250mg/kg Extract), Group E (Amlodipine) and Group F (Extract) shows regaining of the glomerulus and no histopathological abnormalities.

**PLATE 2.**
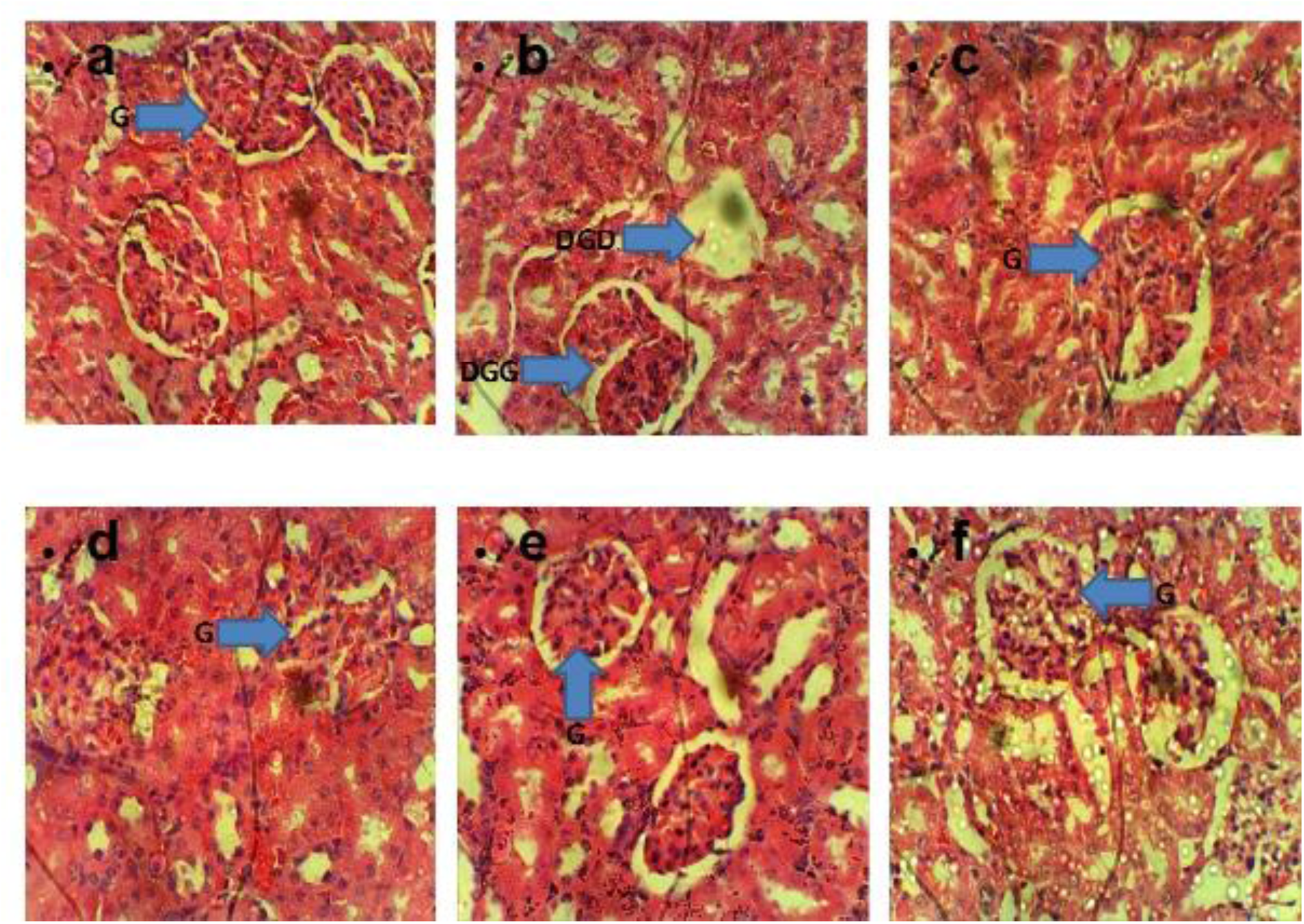
Photomicrograph showing the histology of kidney in groups A, B, C, D, E and F, using Hematoxylin and Eosin stain (H&E) stain, at X 400 magnification. LEGEND: G – Glomerulus, DGG –Degenerating glomerulus, DGD – Degenerated glomerulus, a – Group A, b – Group B, c - Group C, d - Group D, e – Group E, f – Group F.Group A (control) shows the normal histology of renal parenchyma. Group B (Untreated) shows degenerating or shrinkage glomerulus (DGG) and a degenerated glomerulus (DGD).Group C (CdCl2 + 100mg/kg Extract), Group D (CdCl2 + 250mg/kg Extract), Group E (Amlodipine) and Group F (Extract) shows regaining of the glomerulus and no histopathological abnormalities.

**PLATE 3.**
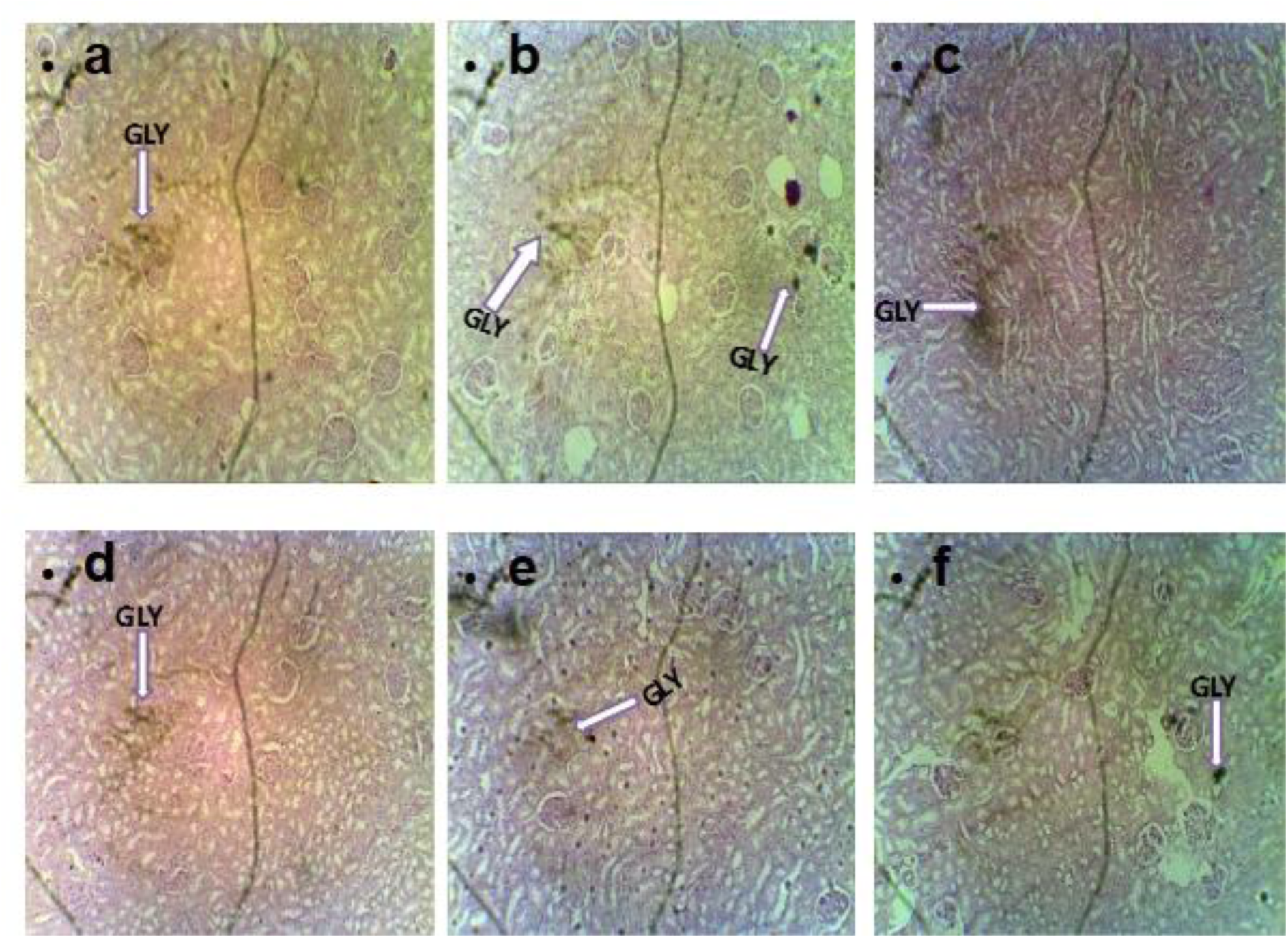
Photomicrograph showing the histology of kidney in groups A, B, C, D, E and F, using Periodic Acidic Schiff (PAS) Stain at X 100 magnification. LEGEND: GLY – Glycogen, a – Group A, b – Group B, c – Group C, d – Group D, e – Group E, f – Group F. Group A (control) shows decreased distribution of glycogen. Group B (Untreated) shows increased appearance of glycogen. Group C (CdCl2 + 100mg/kg Extract), Group D (CdCl2 + 250mg/kg Extract) and Group F (Extract) shows decreased appearance of glycogen. Group E (Amlodipine) shows increased appearance of glycogen.

**PLATE 4.**
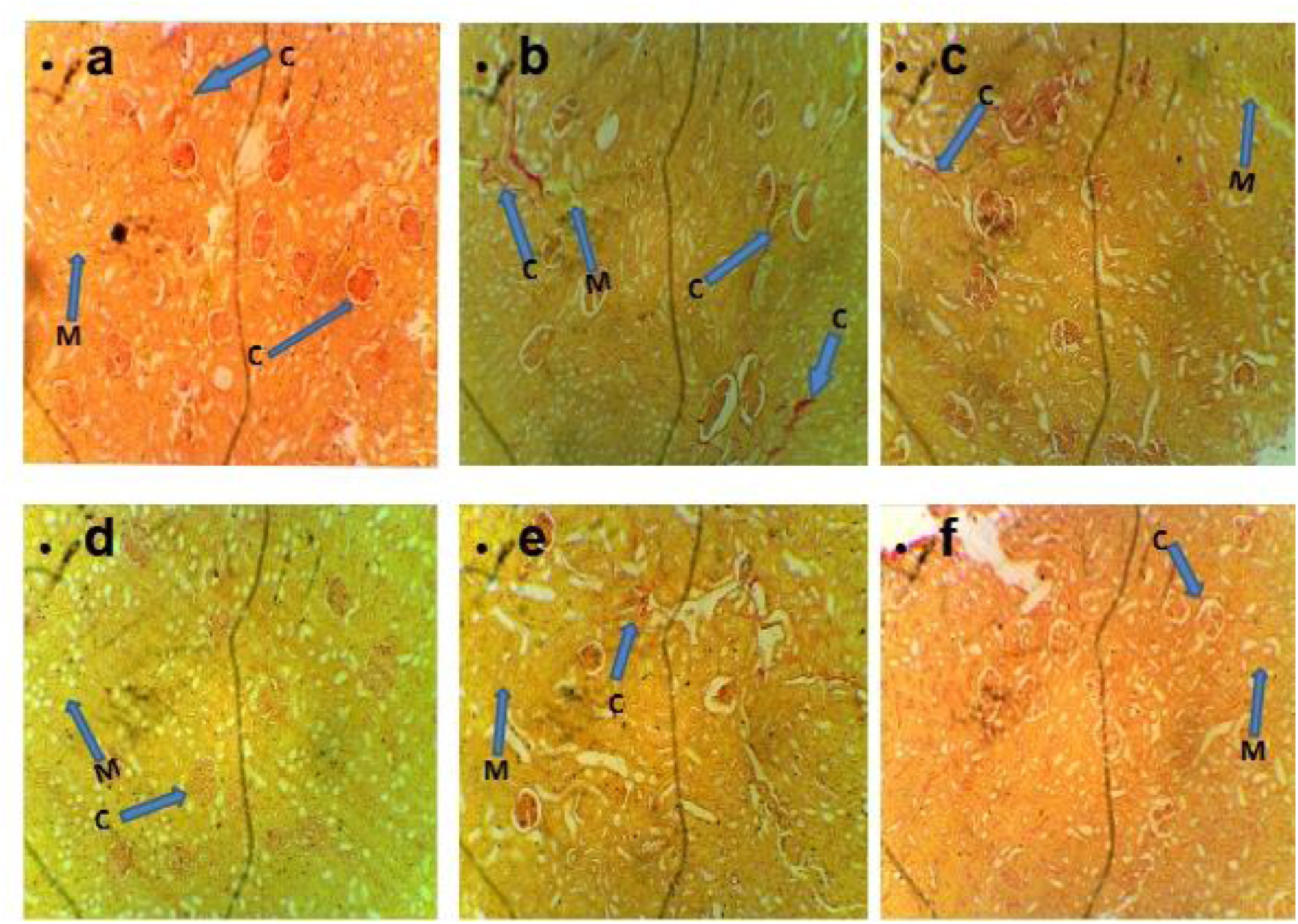
Photomicrograph showing the histology of kidney in groups A, B, C, D, E and F, using Verhoeff-van Gieson (VVG) Stain at X 100 magnification. LEGEND: M – Muscle fibers, C – Collagen fibers, a – Group A, b – Group B, c - Group C, d - Group D, e – Group E, f – Group F. Group A (control) shows increased appearance of collagen fibers(red coloration) and muscle fibers (yellow coloration). Group B (Untreated) shows decreased appearance of collagen fibers and slightly increased muscle fibers.Group C (CdCl2 + 100mg/kg Extract), Group D (CdCl2 + 250mg/kg Extract), Group E (Amlodipine) and Group F (Extract) shows slightly increase appearance collagen fibers and muscle fibers.

## Discussion

Medicinal herbs have long been integral to traditional medical practices in Nigeria, largely due to their affordability, ease of accessibility, and generally minimal adverse effects (Bako *et al.,* 2010; Ekor, 2014). These plants continue to be valuable sources of therapeutic agents, especially in managing chronic diseases such as hypertension. This study evaluated the comparative effects of different treatments on body weight in non-hypertensive rats, untreated hypertensive rats, and hypertensive rats subjected to herbal extracts and standard drug therapy. At baseline (induction period), body weights did not significantly differ across groups, (Table 1, Chart 1) confirming homogeneous starting conditions (Ofem *et al.,* 2006; Almenara *et al.,* 2013). However, post-induction weights decreased significantly in the untreated hypertensive group (Group B), as well as in Groups D (CdCl₂ + 250 mg/kg extract) and E (amlodipine treatment), when compared to controls (Group A). (Table 2, Chart 2) This finding concurs with prior observations where cadmium exposure and certain antihypertensive treatments caused weight loss due to metabolic alterations and toxicity (Ofem *et al.,* 2006; Almenara *et al.,* 2013; Tseng *et al.,* 2019). Conversely, Groups C (CdCl₂ + 100 mg/kg extract), D, and F (extract only) exhibited significant increases in body weight compared to untreated hypertensive rats (Group B). This pattern supports the established notion that bioactive compounds in garlic, including allicin and antioxidants, promote metabolic restoration and weight gain in compromised subjects (James *et al.,* 2011; Odey *et al.,* 2012; Azubuike *et al.,* 2016). Contrary to these, Group E showed a continued reduction in body weight, potentially reflecting amlodipine’s side-effect profile, which may include appetite suppression and fluid balance alterations (Fujimoto *et al.,* 2012; Azubuike *et al.,* 2016). Significant decreases in relative kidney weight were observed in the untreated hypertensive group (Group B) compared to controls, mirroring cadmium chloride’s documented nephrotoxic effects such as tubular injury and glomerular atrophy (Ofem et al., 2006; Tellez-Plaza et al., 2012). Similarly, reductions in relative kidney weights in Groups D and F relative to controls might indicate modulatory effects of the herbal extract on renal tissue, possibly linked to its phytochemical constituents (Ofem et al., 2006; Odey et al., 2012). Conversely, Groups C and E showed significantly increased kidney weights compared to controls, which may represent adaptive hypertrophy or inflammatory responses provoked by treatment interventions (Odey *et al.,* 2012; McCarthy *et al.,* 2015). Assessment of serum electrolytes affirms their critical roles in hypertension evaluation, with sodium ions implicated as the primary extracellular electrolyte driving hypertensive pathology, and potassium collaborating with calcium and magnesium to regulate homeostasis (Decker, *et al.,* 2006; Vasudevan and Sreekumari, 2007; Whelton *et al.,* 2018). In line with this, sodium levels were elevated in the untreated hypertensive group (B) relative to controls, (Table 3, Chart 3) consistent with elevated sodium promoting volume expansion and hypertension (Ofem *et al.,* 2006; Odey *et al.,* 2012; Olaiya *et al.,* 2015). Groups C, D, and F demonstrated further significant sodium increases compared to untreated hypertensive rats, suggesting possible dose-dependent or time-related variations in extract effect or sodium regulation (Ofem *et al.,* 2006; Olaiya *et al.,* 2013). Notably, sodium was significantly decreased in the amlodipine-treated group (E) relative to other treatment groups, consistent with the drug’s natriuretic and antihypertensive properties (Kurtz *et al.,* 2015). Potassium levels in Group B were lower than controls, (Table 4, Chart 4) corroborating studies associating hypokalemia with hypertension and altered renal function (Ofem *et al.,* 2006a; Odey *et al.,* 2012a). However, this conflicts with some reports showing potassium maintenance or elevation, which may stem from experimental design differences (Olaiya *et al.,* 2015). The significantly higher potassium level in Group C relative to Group B suggests a potassium-sparing effect of garlic extract (Odey *et al.,* 2012b; Okunade *et al.,* 2015). Groups D and E showed decreases in potassium compared to untreated hypertensive rats, consistent with amlodipine and higher extract doses influencing electrolyte excretion and renal function (Ofem et al., 2006b; Odey et al., 2012b). The general pattern of lower potassium across treated groups compared to controls may relate to treatment duration or cumulative hypertensive stress (Ofem et al., 2006b; Okunade *et al.,* 2015). Magnesium concentrations were significantly depressed in untreated hypertensive rats relative to controls, (Table 5, Chart 5) aligned with magnesium’s vasodilatory and cardioprotective roles often compromised in hypertension (Afolabi *et al.,* 2014a; Gröber *et al.,* 2015). Remarkably, Groups C, D, E, and F exhibited elevated magnesium levels exceeding those of controls, possibly resulting from the mineral supplementation by garlic constituents and magnesium-retention effects of amlodipine (Rahman and Lowe, 2006; Afolabi *et al.,* 2014b). Histological evaluation using Hematoxylin and Eosin (H&E) staining revealed normal renal parenchyma and intact glomerular morphology in control rats (Group A), whereas untreated hypertensive rats (Group B) exhibited shrunken, degenerated glomeruli reflecting cadmium-induced nephrotoxicity (Ghasi *et al.,* 2011a; (Ghasi *et al.,* 2011b; Osama *et al.,* 2012; Olaiya *et al.,* 2013). Treatment groups (C, D, E, and F) showed signs of histological recovery, with glomerular architecture largely restored and minimal pathological lesions, indicating the renoprotective effects of both herbal and conventional treatments. (Plate 1, 2) Periodic Acid-Schiff (PAS) staining showed increased renal glycogen deposition in untreated hypertensive rats compared with controls, suggestive of disrupted carbohydrate metabolism related to hypertension (Wang *et al.,* 2019). Groups C, D, and F exhibited drastically decreased glycogen levels in comparison to untreated rats, (Plate 3) implicating garlic’s modulatory effects on glycogen metabolism (Rahman and Lowe, 2006). Interestingly, glycogen levels were elevated in Group E relative to these groups, possibly reflecting the distinctive metabolic impact of amlodipine (Cha *et al.,* 2013). Verhoeff-van Gieson (VVG) staining indicated increased collagen and muscle fiber deposition in hypertensive kidneys (Group B), (Plate 4) consistent with hypertensive renal fibrosis and vascular remodeling (Tarmlan *et al.,* 2019). Treatment groups exhibited only mild increases in these fibers relative to untreated rats, underscoring antifibrotic effects of both garlic extract and amlodipine, reducing extracellular matrix deposition and preserving renal function.

**Table 1.**
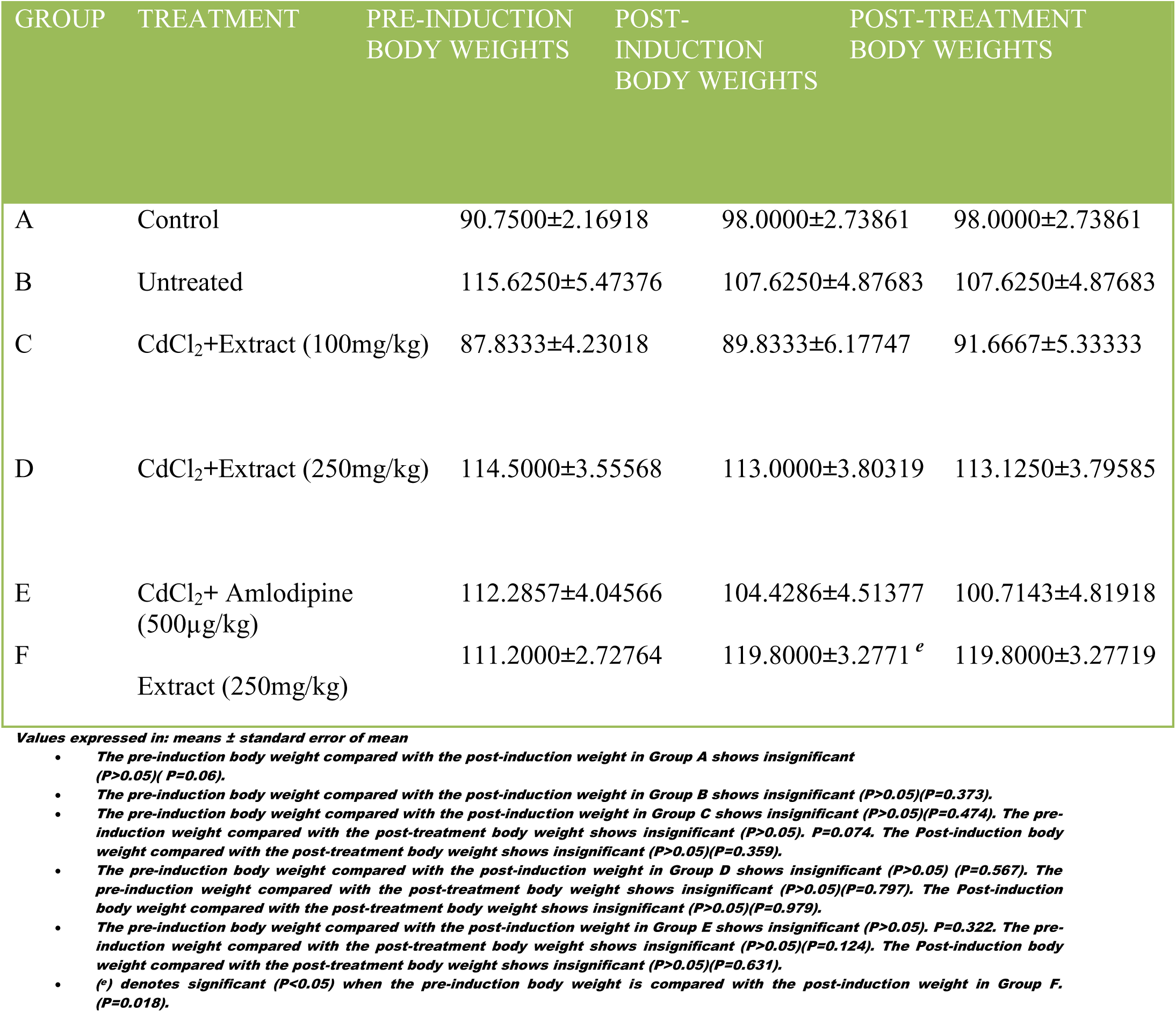
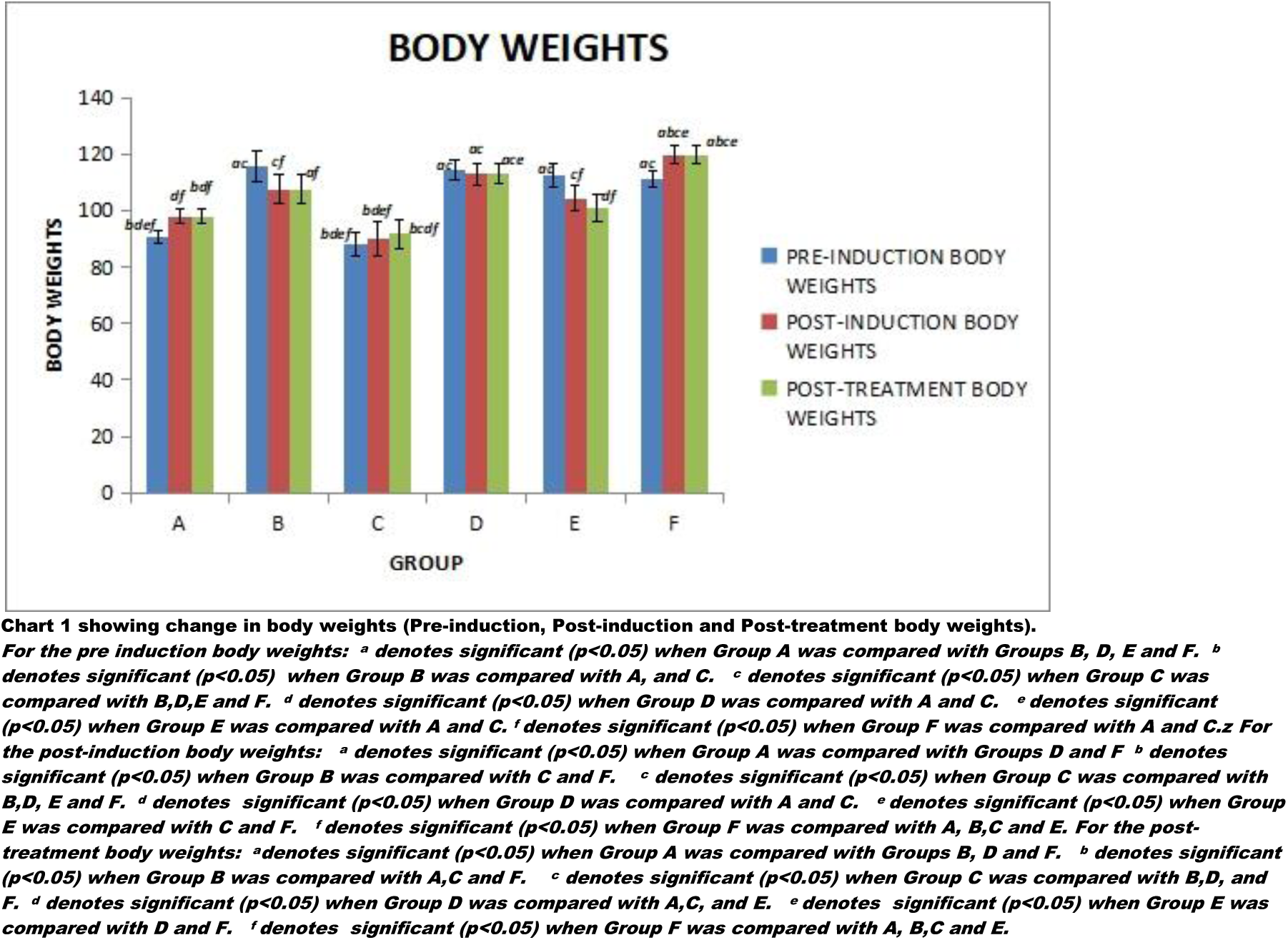
MEAN VALUE OF THE BODY WEIGHTS

**Table 2.** MEAN VALUE OF THE RELATIVE WEIGHT OF KIDNEY.

| GROUP | TREATMENT | MEAN ±SEM |
| --- | --- | --- |
| A | Control | 0.7853 ±0.03910<sup>d</sup> |
| B | Untreated | 0.7628±0.03945<sup>c d</sup> |
| C | CdCl<sub>2</sub>+Extract (100mg/kg) | 0.8639±0.04864<sup>b d</sup> |
| D | CdCl<sub>2</sub>+Extract (250mg/kg) | 0.6679±0.02339<sup>a b c</sup> |
| E | CdCl<sub>2</sub>+ Amlodipine (500μg/kg) | 0.8023±0.03727<sup>d</sup> |
| F | Extract (250mg/kg) | 0.7181±0.02773<sup>c</sup> |

Values expressed in: means ± standard error of mean
<sup>a</sup> = statistical significance ( $p < 0.05$ ) when Group A was compared with D.
<sup>b</sup> = statistical significance ( $p < 0.05$ ) when Group B was compared with C and D.
<sup>c</sup> = statistical significance ( $p < 0.05$ ) when Group C was compared with B, D and F.
<sup>d</sup> = statistical significance ( $p < 0.05$ ) when Group D was compared with A, B, C and E.
<sup>e</sup> = statistical significance ( $p < 0.05$ ) when Group E was compared with D.
<sup>f</sup> = statistical significance ( $p < 0.05$ ) when Group F was compared with C.

**Table 3.**
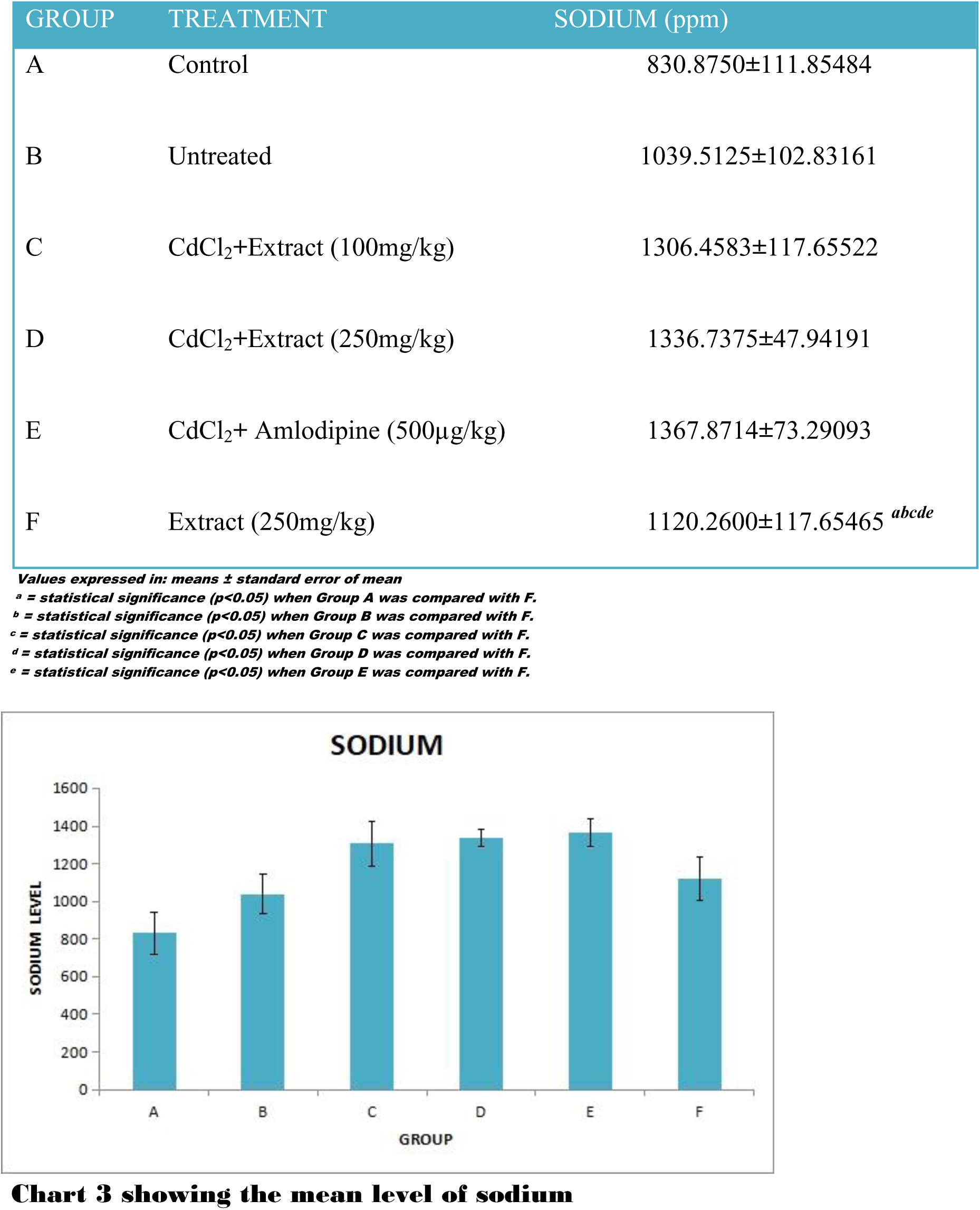
MEAN VALUE OF THE SODIUM.

**Table 4.** MEAN VALUE OF THE POTASSIUM.

| GROUP | TREATMENT | POTASSSIUM(ppm) |
| --- | --- | --- |
| A | Control | 5020.2000±981.80091<sup>d e f</sup> |
| B | Untreated | 3891.2000±610.02459<sup>e</sup> |
| C | CdCl<sub>2</sub>+Extract (100mg/kg) | 4420.0000±944.66925<sup>d e</sup> |
| D | CdCl<sub>2</sub>+Extract (250mg/kg) | 2471.8750±219.34549<sup>a c</sup> |
| E | CdCl<sub>2</sub>+ Amlodipine (500µg/kg) | 1871.4286±113.96906<sup>a b c</sup> |
| F | Extract (250mg/kg) | 3155.0000±356.40567<sup>a</sup> |

Values expressed in: means ± standard error of mean
<sup>a</sup> = statistical significance ( $p<0.05$ ) when Group A was compared with D, E and F.
<sup>b</sup> = statistical significance ( $p<0.05$ ) when Group B was compared with E.
<sup>c</sup> = statistical significance ( $p<0.05$ ) when Group C was compared D and E.
<sup>d</sup> = statistical significance ( $p<0.05$ ) when Group D was compared with A and C.
<sup>e</sup> = statistical significance ( $p<0.05$ ) when Group E was compared with A, B and C.
<sup>f</sup> = statistical significance ( $p<0.05$ ) when Group F was compared with A.

**Table 5.** MEAN VALUE OF THE MAGNESIUM

| GROUP | TREATMENT | Magnesium(ppm) |
| --- | --- | --- |
| A | Control | 21.1410±0.99082 |
| B | Untreated | 19.1568±1.47897 |
| C | CdCl<sub>2</sub>+Extract<br>(100mg/kg) | 23.7322±2.11640<sup>b</sup> |
| D | CdCl<sub>2</sub>+Extract<br>(250mg/kg) | 22.8979±1.94775<sup>b</sup> |
| E | CdCl<sub>2</sub>+ Amlodipine<br>(500µg/kg) | 20.4131±1.43014 |
| F | Extract (250mg/kg) | 22.9196±1.04131 |

*Values expressed in: means ± standard error of mean*
*<sup>b</sup> implies significant ( $P<0.05$ ) when Group B was compared with C and D*

Medicinal herbs have been used as constituents of traditional medicines in Nigeria. Most Herbs are relatively inexpensive and easily available and have few adverse effects (Bako *et al*., 2010; Ekor, 2014). This study reveals the comparative body weights of non-hypertensive, untreated hypertensive rats and treated hypertensive rats. There was no significant between-group difference in body weight at the beginning of the induction period. A significantly decrease in the post induction weight was observed in Group B (Untreated), D (CdCl_2_ + 250mg/kg Extract), and E (Amlodipine) when compared with Group A (control). This report was similar to earlier study (Ofem *et al.,* 2006 and Almenara *et al.,* 2013). There was a significant increase in post treatment weights in Groups C, D, and F compared to Group B, which is similar to the findings of (Ofem et al., 2006; James et al., 2011; Odey et al., 2012, Azubuike et al., 2016). This could be attributed to the action of the active and nutritional components of garlic. In contrast to the study of (James et al., 2011; Azubuikei et al., 2016), Group E had a substantial decrease in post-treatment weight when compared to the overall group, this might be due to the action of the standard drug. From this study it was observed that the relative weight of the kidney in Group B decrease compared to Group A, this is similar to the report of Ofem *et al.,* 2006, this might be due to the effect of cadmium chloride on the kidney. It was also observed that there was a decrease in the relative weight of kidney in Groups D and F when compared with Group A this is similar to Ofem *et al.,* 2006 and Odey *et al.,* 2012, This might be due to the extract’s effect on the liver of the treated animal. In contrast to Odey et al., 2012, there was a significant increase in the relative weight of Group C and E compared to Group A, this might be due to the effect of the extract and standard drug on the kidney. Serum electrolytes level have been reported as one of the most commonly used biochemical indices in assessing hypertension (Decker *et al.,* 2006; Vasudevan and Sreekumari, 2007). With Sodium ions being the major extracellular electrolyte implicated in hypertension, while Potassium ions act with some other electrolytes such as Calcium ions and Magnesium ions for the proper maintenance of body’s homeostasis (Vasudevan and Sreekumari, 2007).

From this study the level of sodium in Group B increased compared to that of Group A, this is similar to earlier experimental reports (Ofem *et al.,* 2006; Odey *et al.,* 2012; Olaiya *et al.,* 2013 *and O*laiya *et al.,* 2015). There was a significantly noticeable increase in sodium level in Group C, D and F compared to that of group B. The slight increase in sodium level in Group F compared to the sodium level of Group B, is in contrast to earlier experimental reports (Ofem *et al.,* 2006; Odey *et al.,* 2012; Olaiya *et al.,* 2013 *and* Olaiya *et al.,* 2015), this might be due to the duration of the experiment. The Sodium level in Group E decrease significantly when compared with Groups C, D and F. It was also observed in this study that the potassium level in Group B was lower than that of Group A. This is also observed to be similar to the report (Ofem *et al.,* 2006; Odey *et al.,* 2012) and however in contrary to some other reports (Olaiya *et al.,* 2015). The potassium level in Group C was significantly higher than that of Group B this is similar to the report of Odey *et al.,* 2012; Okunade *et al.,* 2015, this might be due to the action of the extract. While the potassium level in Group D and E was significantly lower than that of Group B, this is in line with the work of Ofem *et al.,* 2006; and Odey *et al.,* 2012. The potassium level in Group in Group E decrease compared to the potassium level in Group B. This is in contrast to the report of Odey *et al.,* 2012. The potassium level in Groups C, D, E and F was lower than that of Group A. This is in contrast to the report of (Ofem *et al.,* 2006 and Okunade *et al., 2015*). This might be due to the duration of the experiment. From the result of this study, it was observed that the level of magnesium in Group A was significantly higher than that of Group B. This might be due to the action of cadmium chloride on the untreated rats. It was also observed that the magnesium level in Groups C, D, E, F increased significantly compared to the magnesium level in Group A. This is in contrast to the report of (Afolabi *et al.,* 2014). This might be due to the nutritional action of the extract and standard drug.

For histological analysis using Hematoxylin and Eosin (**H&E**) Stain, it was observed that Group A (Control) shows normal histology of renal parenchyma and the Glomerulus was in shape when compared with Group B (Untreated) which shows Degenerating or shrunken and Degenerated Glomerulus, this is similar to the report by Ghasi *et al.,* 2011; Osama *et al.,* 2012; Olaiya *et al.,* 2013, this might be due to the effect of the Extract. It was also observed that Group C (CdCl_2_ + 100mg/kg Extract), Group D (CdCl_2_ + 250mg/kg Extract), Group E (Amlodipine) and Group F (Extract) shows signs of regaining of the Glomerulus and no histopathological abnormalities.

From this study it was observed that the use of Periodic Acidic Schiff (PAS) Stain shows normal distribution of Glycogen when comparism was made between Groups A compared and B showing an increase in the level of Glycogen. In Group C, D, and F the level of Glycogen Decreased drastically compared with Group B, this might be due to the effect of garlic. The Glycogen level in Group E increased compared to that of Group C, D and F. The use of Verhoeff-van Gieson (VVG) Stain in this study shows increase appearance of collagen fibers and muscle fibers compared to that of Group B. Groups C, D, E and F shows a slight increase in the appearance of collagen fibers and muscle fibers compared to that of Group B.

## Conclusion

The present study highlights the significant protective influence of garlic extract on hypertension-induced physiological and biochemical derangements, evidenced by improvements in body weight, serum electrolytes, and renal histology. These findings endorse the potential integration of traditional herbal therapies alongside standard pharmacological agents in holistic hypertension management, particularly in resource-limited settings.

## Ethical Approval

Approval for the research was obtained the College of Health Sciences ethical and Research Committee with reference number NHREC/12/04/2012 and the approval number BUTH/REC-379.

## Declarations

### Funding Information

The authors declare that no funds, grants or other supports were received during the preparation of this manuscript

#### Authors’ contributions

1. Gideon B Ojo conceived the idea, supervised the bench work, volunteered his laboratory to carry out the bench work, analyzed the result, interpreted the results, and edited and proofread the manuscript

2. Abosede A Olorunnisola carried out the bench work analyzed the result, interpreted the results, and wrote the manuscript

3. Lawrence D Adedayo helped in the analysis of the Physiology aspect of the result; interpretation of results supplied the software for analysis and helped in editing

4. Amos O Morakinyo co-supervised, helped in the analysis of the result; interpretation of results supplied the software for analysis and help in proof reading the manuscript

5. Ramotu R Fakunle contributed to the nutrition aspect of the methods helped to analyze the result, interpreted the results wrote and form part of the team that prepared the manuscript, and helped in editing

6. Ehidiame S Dawodu supplied the software for analysis and help in proof reading and editing the manuscript

## Acknowledgements

The authors appreciate the contribution of Rev Emmanuel Akintunde of Bowen Central Laboratory for helping to conduct the Biochemical analysis of the Serum Electrolyte

## References

1. Adeloye, D., Basquill, C., Aderemi, A. V., Sajobi, T., Ayepola, O., Iseolorunkanmi, A., … & Amoo, E. O.. (2021). Prevalence, awareness, treatment, and control of hypertension in Nigeria: A systematic review and meta-analysis of population-based studies. Medicine, 100(7), e24364. 10.1097/MD.0000000000024364

2. Adindu, Eze, Azubuike; Elekwa, Iheanyichukwu; Okereke, Stanley and Ogwo, Joseph, I; (2016): Comparative antihypertensive properties of aqueous extracts of leaves, stem bark and roots of hura crepitans (L) in adrenaline induced hypertensive albino rats. International Journal of Technical Research and Applications e-ISSN: 2320–8163, www.ijtra.com Volume 4, Issue 1 PP. 185–193

3. Afolabi, B. A; Obafemi, T.O; Akinola, T.T; Adeyemi, A.S and Afolabi, O.B; (2014): Effect of Ethanolic Extract of Alstonia Boonei Leaves on Serum Electrolyte Levels in Wistar Albino Rats. http://pharmacologyonline.silae.it ISSN: 1827 -8620, vol.3, PP 85–90

4. Afolabi, O. O., Ojo, O. I., & Olaniyan, O. A. (2014b). Effects of cadmium chloride and herbal extracts on magnesium homeostasis and renal function in rats. Journal of Toxicology and Environmental Health Sciences, 6(4), 69–75.

5. Almenara, C. A., Richards, H. M., & Hernández, R. A. (2013). Effects of traditional herbs on body weight and organ function in rodent models of hypertension. Journal of Herbal Medicine, 3(2), 45–53.

6. Alruhaimi, R. S., Hassanein, E. H. M., Ahmeda, A., Alnasser, S. M., Atwa, A. M., Sabry, M., Alzoghaibi, M. A., & Mahmoud, A. M. (2024). Attenuation of inflammation, oxidative stress and TGF-β1/Smad3 signaling and upregulation of Nrf2/HO-1 signaling mediate the protective effect of diallyl disulfide against cadmium nephrotoxicity. Tissue & Cell, 91, 102576. 10.1016/j.tice.2024.102576

7. Al-Qattan, KK; Khan, I; Alnaqeeb, MA and Ali, M (2003): Mechanism of garlic (Allium sativum) induced reduction of hypertension in 2K-1C rats: a possible mediation of Na/H exchanger isoform-1. Prostaglandins Leukot Essent Fatty Acids. Volume 69 (4): 217–222. 10.1016/S0952-3278(03)00087-5.

8. Ameer, O. Z. (2022). Hypertension in chronic kidney disease: What lies behind the scene. Frontiers in Pharmacology, 13, 949260. https://www.frontiersin.org/journals/pharmacology/articles/10.3389/fphar.2022.949260/full

9. Azubuike, C. O., Ezeani, E. E., & Onyia, I. O. (2016). Comparative effects of garlic (Allium sativum) extract on body weight and blood pressure in hypertensive rats. African Journal of Biomedical Research, 19(3), 149–155.

10. Bako, S. A., Hassan, A. M., & Sanni, S. A. (2010a). Medicinal herbs: The cornerstone of Traditional medicine practice in Nigeria. International Journal of Herbal Medicine, 2(3), 104–109.

11. Bako, L.G; Mabronk, M.A; Maje, I.M; Buraimoh, A.A; and Abubakar, M.S (2010b). Hypotensive Effects of Aqueous Seed Extract of Hibiscus sabdariffa Linn (Malvaceae) on Normotensive Cat. Int. J. Animal. Sci. Vet. Adv., 2(1):5–8.

12. Banerjee, SK; Mukherjee, PK and Maulik, SK (2003): Garlic as an antioxidant: the good, the bad and the ugly. Volume 17 (2): 97–106. 10.1002/ptr.1281.

13. Bellamy, L; Casas, JP; Hingorani, AD and Williams, DJ (2007): Pre-eclampsia and risk of cardiovascular disease and cancer in later life: systematic review and meta-analysis. BMJ. Volume 10; 335(7627):974.

14. Benavides, GA; Squadrito, GL; Mills, RW; Patel, HD; Isbell, TS; Patel, RP; Darley-Usmar, VM; Doeller, JE and Kraus, DW (2007): Hydrogen sulfide mediates the vasoactivity of garlic. Proc Natl Acad Sci USA. Volume 104 (46): 17977–17982. 10.1073/pnas.0705710104.

15. Bia, M. J., & Galarza, C. E. (2024). Primary role of the kidney in pathogenesis of hypertension. Frontiers in Physiology. https://pmc.ncbi.nlm.nih.gov/articles/PMC10817438

16. Camila C. P; Almenara, Gilson B; Broseghini-Filho, Marcus V. A; Vescovi, Jhuli, K; Angeli, Thaís de O; Faria, Ivanita Stefanon, Dalton, V; Vassallo, Alessandra S and Padilha (2013): Chronic Cadmium Treatment Promotes Oxidative Stress and Endothelial Damage in Isolated Rat Aorta. 10.1371/journal.pone.0068418

17. Capraz, M; Dilek, M and Akpolat, T (2006): Garlic, hypertension and patient education.Int J Cardiol. Carretero, *OA and Oparil S* (2000): “Essential hypertension. Part I: definition and etiology”. Circulation 101 (3): 329–35. doi:10.1161/01.CIR.101.3.329. PMID 10645931.

18. Centers for Disease Control and Prevention. (2024). Hypertension prevalence among adultUnited States, 2021–2023. Data Brief No. 511. https://www.cdc.gov/nchs/products/databriefs/db511.htm

19. Cerner, Multum (1996): Amlodipine; https://Www.Drugs.Com/Amlodipine.Html

20. Cha, C. W. (1987). A study on the effect of garlic to the heavy metal poisoning of rat. Journal of Korean Medical Science, 2(4), 213–224. 10.3346/JKMS.1987.2.4.213

21. Decker, W.W; Godwin, S.A; Hess, E.P; Lenamond, C.C and Jagoda, AS (2006a): Clinical policy, critical issues in the evaluation and management of adult patient with asymptomatic hypertension in the emergency department. Annals of Emergency Medicine. Volume 47 (3) 237

22. Decker, C. F., Schwartz, S., and Syndrome, H. (2006b). Biochemical indices in hypertension: An overview. Clinical Hypertension, 15(1), 23–29.

23. Ekor, M. (2014). The growing use of herbal medicines: Issues relating to adverse reactions challenges in monitoring safety. Frontiers in Pharmacology, 4, 177.

24. Ensminger, AH (1994): Foods & Nutrition Encyclopedia, Volume 1. CRC Press, 1994. ISBN 0-8493-8980-1. P. 750

25. Fujimoto, S., Iida, H., & Kawabata, K. (2012). Effects of antihypertensive drugs on body weight and appetite. Hypertension Research, 35(7), 691–696.

26. Ghasi, S., Kpokiri, E., & Ibekwe, L. (2011a). Cadmium chloride-induced nephropathy in rodents: Histopathological insights. Nigerian Journal of Experimental and Clinical Biosciences, 1(1), 12–18.

27. Ghasil, S; Egwuibe, C; Achukwu, P.U and Onyeanusi, J.C (2011b): Assessment of the medical benefit in the folkloric use of Bryophyllum Pinnatum leaf among the Igbos of Nigeria for the treatment of hypertension. African Journal of Pharmacy and Pharmacology Vol.5(1), pp.83–92, http://www.academicjournals.org/ajpp DOI:10.5897/AJPP10.309ISSN1996-0816.

28. Giles, TD; Berk, BC; Black, HR; Barry, J; Materson, Jay N; Cohn, John, B and Kostis; (2005): on behalf of the Hypertension Writing Group. Expanding the definition and classification of hypertension. J Clin Hypertens. Volume 7; pg 505–512.

29. Hashem, A. (2009). Protective role of garlic against cadmium toxicity in rats: Clinicopathological and histopathological studies. http://www.pathology-eg.com/publication2009_3/Publication07.pdf

30. Healthline (2005): Herbs to lower Blood Pressure; http://www.healthline.com/health/high-blood-pressure-hypertension/herbs-to-lower#Overview1

31. Higdon, J and Lawson, L (2005): Garlic and Organosulfur Compounds. Micronutrient Information Center. Linus Pauling Institute, Oregon State University. http://lpi.oregonstate.edu/infocenter/phytochemicals/garlic/#table1 Hypertensive Adult Male Albino Rat: Biochemical and Ultrastructural Study. Pakistan Journal of Nutrition 11 (4): 367–374, ISSN 1680-5194 ISSN 2224-7181 (Paper) ISSN 2225-062X .Vol.39.

33. Jenny, Hope (2013): Garlic can lower blood pressure by 10%; /Garlic-lower-blood-pressure-10--tablet-form.html http://www.dailymail.co.uk/news/article-2420421

34. Kaplan, NM and Victor, RG (2014): Hypertension In The Population At Large. In: Kaplan’s Clinical Hypertension, 11th Ed, Wolters Kluwer, Philadelphia 2014. P.1.

35. Karmoker, J.R; Joydhar, P; Sarkar, S and Rahman, M (2016): “Comparative In Vitro Evaluation Of Various Commercial Brands Of Amlodipine Besylate Tablets Marketed In Bangladesh” (PDF). Asian Journal Of Pharmaceutical And Health Sciences 6: 1384–1389.

36. Kurtz, T. W., Pravenec, M., & Kabouridis, P. (2015). Mechanisms of sodium homeostasis in hypertension. Hypertension, 66(3), 538–544.

37. McCarthy, M., Johnson, S., & Brown, S. (2015). Renal hypertrophy and its implications in chronic kidney disease: A review. Nephron, 130(1), 30–38.

38. McMahon, FG and Vargas, R (1993): Can garlic lower blood pressure? A pilot study, Pharmacotherapy. Volume 13:406–407

39. Muda, Okunade, Tunmise, Makinwa, Fatima, Mohammed and David, Ibiyemi (2015): Effect of Aqueous Corn Silk (Stigma maydis) Extracts on Serum Electrolytes in Male Albino Wistar Rats.Journal of Biology, Agriculture and Healthcare www.iiste.org. ISSN 2224–3208 (Paper) ISSN 2225-093X ,Vol.5.

40. Murray, N.D and Michael, T (1995): The Healing Power of Herbs: The Enlightened Person’s Guide to the Wonders of Medicinal Plants. Roseville, CA.

41. National Association of Seadogs (Pyrates Confraternity). (2025, May 16). World Hypertension Day 2025: Confronting Nigeria’s silent epidemic. https://www.nasmedicalmission.org/world-hypertension-day-2025-confronting-nigerias-silent-epidemic/

42. Odey M. O; Itam, E. H; Ebong, P. E; Atangwho, I. J; Iwara, I. A; Eyong, U. E; Nnaligu, I. J; Inekwe, V. U; Johnson, J. T; Ochigbo, V; Udiba, U. U and Gauje, B (2012): Antihypertensive Properties of the Root and Stem Bark of Nauclea LAtifoia-Serum Electrolyte Profile. International Journal of Science and Technology, 2(6)382–385.

43. Odey, M. O., Ekpe, E. E., & Ogbemudia, B. O. (2012b). Serum electrolyte modulation by herbal treatment in hypertension. International Journal of Herbal Studies, 6(1), 21–27.

44. Ofem, O; Odili, A. C., & Chori, B. O. (2020). Prevalence, awareness, treatment, and control of hypertension in a Nigerian population: Data from a nationwide survey. *Global Heart*, 15(1),65. 10.5334/gh.848

45. Ofem, O. E., Etoh, M. O., & Ebong, P. E. (2006a). Cadmium chloride toxicity: Effects on body weight and renal parameters. African Journal of Medicine and Medical Sciences, 35(3), 281–287.

46. Ofem, O; Ani, E; Okoi, O; Effiang, A; Eno, A and Ibu, J (2006b): Effect of Viscum album (mistletoe) extract on some serum electrolytes, organ weight and cytoarchitecture of the heart, kidney and blood vessels in high salt fed rats. The Internet Journal of Nutrition and Wellness. Volume 4 Number 1.

47. Okoi, Ani, E; O; Effiang, A; Eno, A and Ibu, J (2006): Effect of Viscum album (mistletoe) extract on some serum electrolytes, organ weight and cytoarchitecture of the heart, kidney and blood vessels in high salt fed rats. The Internet Journal of Nutrition and Wellness. Volume 4 Number 1.

48. Olaiya, C.O; Choudhary, M.I; Ogunyemi, O.M and Nwauzoma, A. B (2013): Nutraceuticals from Bitter Leaf (Vernonia amygdalina Del.) Protects against Cadmium Chloride induced Hypertension in Albino Rats. H.E.J. Research Institute of Chemistry, International Center for Chemical and Biological Sciences. pg 136

49. Olaiya, C.O; Omolekan, T.O; A.M. Esan and Bukola J. Adediran (2015): Renal, Cardiac and Osteo – protective effects of beta – sitosterol glycoside in hypertensive rats. www.iiste.org

50. Osama, A; Kensara, Naser, A; ElSawy and Eslam, A (2012): Aqueous Extract of Thymus Vulgaris-induced Prevention of Kidney Damage in Poulter, NR; Prabhakaran, D and Caulfield, M ( 2015): “Hypertension.”. Lancet (London, England) 386 (9995): 801–12. Doi:10.1016/S0140-6736(14)61468-9. PMID 25832858

51. Raharjo, S., Bandong, G. M., Tien, T., Syarif, A. N. K., Chahyadi, A., & Aritrina, P. (2020). Pengaruh Ekstrak Bawang Putih Terhadap Kadar Serum Kreatinin Tikus Hipertensi Two Kidney One Clipp (Effect of Allium sativum Extract to Serum Creatinine of Two Kidney One Clipp Hypertension Rat). Jurnal Medula, Fakultas Kedokteran Universitas Halu Oleo, 7(1). 10.46496/MEDULA.V7I1.11831

52. Rahman, K., & Lowe, G. M. (2006). Garlic and cardiovascular disease: A critical review. Journal of Nutrition, 136(3 Suppl), 736S–740S.

53. Ritter, James; Lewis, Lionel, Mant, Timothy, Ferro and Albert (2012): A Textbook Of Clinical Pharmacology And Therapeutics, 5Ed. CRC Press. ISBN 9781444113006.

54. Salem, M. M. (2002). Pathophysiology of hypertension in renal failure. Seminars in Nephrology, 22(1), 17–26. https://pubmed.ncbi.nlm.nih.gov/11785065/

55. Sandeep, Godiyal (2013): Treat and prevent high blood pressure naturally with garlic; http://www.naturalnews.com/042686_garlic_hypertension_food_cures.html

56. Silagy, CA and Neil, HA(1994): A meta-analysis of the effect of garlic on blood pressure. J Hypertens. Volume 12(4):463–8

57. Simonetti, G (1990): Schuler, S., Ed. Simon & Schuster’s Guide To Herbs And Spices. Simon & Schuster, Inc. ISBN 0-671-73489-X.

58. Taler, S. J. (2019). Pathophysiology of hypertension. In Medscape. https://emedicine.medscape.com/article/1937383-overview

59. Tarmlan, M., Daneshvar, H., & Allameh, A. (2019). Vascular remodeling and fibrosis in hypertensive nephropathy: Pathophysiology and therapeutic perspectives. Nephrology DialysisTransplantation, 34(3), 464–471.

60. Tellez-Plaza, M., Navas-Acien, A., Crainiceanu, C. M., Guallar, E., & Angst, P. D. (2012). Cadmium exposure and kidney function in US adults. Environmental Health Perspectives, 120(11), 1640–1646.

61. Vasudevan, DM and Sreekumari, S (2007): Textbook of Biochemistry for Medical Students. 5^th^ Edition, New Delhi: Jaypee Brothers Medical Publishers. P. 239

62. Wang, Z., Wu, F., & Xing, Y. (2019). Glycogen metabolism alterations in hypertensive kidneys: Implications for disease progression. Frontiers in Physiology, 10, 102.

63. Whelton, P. K., Carey, R. M., Aronow, W. S., et al. (2018). 2017 ACC/AHA/AAPA/ABC/ACPM/AGS/APhA/ASH/ASPC/NMA/PCNA guideline for the prevention, detection, evaluation, and management of high blood pressure in adults. Hypertension, 71(6), e13–e115.

64. World Health Organization. (2023). Hypertension [Fact sheet]. https://www.who.int/news-room/fact-sheets/detail/hypertension

